# Analysis of *DNM3* and *VAMP4* as genetic modifiers of *LRRK2* Parkinson’s disease

**DOI:** 10.1101/686550

**Authors:** EE Brown, C Blauwendraat, J Trinh, M Rizig, MA Nalls, E Leveille, JA Ruskey, H Jonvik, MMX Tan, S Bandres-Ciga, S Hassin-Baer, K Brockmann, J Infante, E Tolosa, M Ezquerra, S Benromdhan, M Benmahdjoub, J Hardy, AB Singleton, RN Alcalay, T Gasser, D Grosset, NM Williams, A Pittman, Z Gan-Or, R Fernandez-Santiago, A Brice, S Lesage, M Farrer, N Wood, HR Morris, on behalf of the International Parkinson Disease Genomics Consortium (IPDGC)

## Abstract

**Objective:** To assess genetic modifiers of Parkinson’s disease (PD) age at onset (AAO) penetrance in individuals carrying common and rare *LRRK2* risk alleles

**Methods:** We analysed reported genetic modifier *DNM3* rs2421947 in 724 *LRRK2* p.G2019S heterozygotes using linear regression of AAO. We meta-analysed our data with previously published data (n=754). *VAMP4* is in close proximity to *DNM3* and is associated with PD. We analysed the effect of the rs11578699 *VAMP4* variant on pG2019S penetrance in 786 *LRRK2* p.G2019S heterozygotes. We also evaluated the impact of *VAMP4* variants using AAO regression in 4882 patients with PD carrying a common *LRRK2* risk variant (rs10878226).

**Results:** There was no evidence for linkage disequilibrium between DNM3 rs2421947 and *VAMP4* rs11578699. Our linear regression AAO of 724 p.G2019S carriers showed no relationship between *DNM3* rs2421947 and AAO (beta = −1.19, p = 0.55, n =708). Meta-analysis with previously published data did not indicate a significant effect on AAO (beta = −2.21, p = 0.083, n = 1304), but there was significant heterogeneity in the analyses of new and previously published data. *VAMP4* rs11578699 was nominally associated with AAO in patients dichotomized by the common *LRRK2* risk variant rs10878226 (beta=1.68, se=0.81 p=0.037).

**Interpretation:** Analysis of *DNM3* in previously unpublished data does not show an interaction between *DNM3* and *LRRK2* G2019S for AAO, however the inter-study heterogeneity may indicate ethnic-specific effects of *DNM3* rs2421947. Analysis of sporadic PD patients stratified by the PD risk variant rs10878226 indicates a possible interaction between *LRRK2* and *VAMP4*.

## Introduction

The p.G2019S coding variant in the *LRRK2* gene is the most common high penetrance mutation causing parkinsonism. The mutation occurs in 1-40% of PD cases, varying by ethnicity (1). *LRRK2* parkinsonism is broadly similar to “idiopathic” disease in clinical manifestations and age at onset (AAO), generating interest in its potential as a therapeutic target with broader application to PD (2). In addition to rare pathogenic mutations, genome wide association studies (GWAS) show that common variation in *LRRK2* is a risk factor for sporadic PD (3) (23).

There is a wide range in PD AAO among p.G2019S carriers. It is estimated that the heritability of PD AAO (~0.11) is lower than the overall heritability of risk for PD (~0.27) (4). Currently the strongest genetic association with AAO is through genetic risk score (GRS). There is overlap between the heritability of risk, as quantified in the GRS, and AAO. As *LRRK2* is a functionally complex protein with many protein-protein interaction domains, there may be diverse genetic modifiers that vary in effect size and frequency. Trinh and colleagues (5) presented evidence of modification by a common variant haplotype tag in *DNM3* (rs2421947), for which the median onset of *LRRK2* parkinsonism for GG homozygotes was 12·5 years younger than that of CC homozygotes (direction of effect GG < GC < CC), suggesting that the C allele or a linked variant may be protective against the development of PD. Similarly, Fernandez-Santiago and colleagues (6) observed a consistent trend for the rs2421947 G allele on AAO in *LRRK2* p.G2019S carriers, with median onset 3 years younger in patients with the G allele, though the result was not significant. Foo and colleagues (2018) found no impact of *DNM3* in PD AAO in individuals carrying Asian *LRRK2* risk alleles (24).

In the largest GWAS meta-analysis of idiopathic PD the nearby *VAMP4* gene variant rs11578699, located 113,325 bp from *DNM3* rs2421947, emerged as a statistically significant PD risk association, raising the possibility the effect of *DNM3* may relate to linkage disequilibrium with *VAMP4* (7). *LRRK2* has a role in both monogenic and idiopathic/sporadic forms of PD and overlap in genetic modifiers may exist.

We carried out a large replication candidate gene study investigating the impact of reported genetic modifier *DNM3* rs2421947 and the nearby GWAS association *VAMP4* rs11578699 on AAO. We specifically sought to replicate the discovery finding, to determine whether the association might vary with ethnicity and whether linkage disequilibrium with VAMP4 may be relevant to this association. We analyzed a multi-ethnic cohort of European, North African, and Ashkenazi Jewish *LRRK2* p.G2019S carriers and a larger cohort of European patients with and without the *LRRK2* common risk allele.

## Methods

### Data collection

#### Patient cohorts

*LRRK2* p.G2019S heterozygotes were identified from cohorts in the International Parkinson’s Disease Genomics Consortium (IPDGC) and other collaborative centres. Data from 724 *LRRK2* p.G2019S heterozygotes was contributed by the National Institute of Health (NIH), University College London (UCL), McGill University (MU) (samples were collected in Columbia University (CU) and Sheba Medical Centre (SMC)), Sorbonne University (SU) (in collaboration with Habib Bourguiba hospital, Sfax, Tunisia, and Blida Hospital, Blida, Algeria), National Institute of Neurological Disorders and Stroke (NINDS), and the University of T**ü**bingen (TU). Data from two published studies analyzing *DNM3* rs2421947 and *LRRK2* p.G2019S parkinsonism with AAO was also included comprising 754 participants (5, 6). All studies willing to participate and currently holding minimum required data for *LRRK2* p.G2019S carriers were included: PD onset age or sampling age for asymptomatic carriers; and *DNM3* rs2421947 or a high RSQ proxy SNP. Where these studies had also genotyped *VAMP4* rs11578699 or a high RSQ proxy, these data were also included in the analysis of the *VAMP4* gene (n=786). Patients with PD without a known Mendelian or high risk genetic cause of disease were collated from the UK-wide *Tracking Parkinson’s* study (8) and from IPDGC datasets. Summary statistics for *DNM3* rs2421947 and *VAMP4* rs11578699 were obtained from the most recent large-scale PD GWAS (7) and PD AAO GWAS (4). All studies were approved by each respective institutional ethics review committee and all participants provided written informed consent. All studies were carried out in accordance with the Declaration of Helsinki (9).

#### Genetic analyses

Genotyping was performed on different platforms (Neurochip array, NeuroX array, TaqMan assays, Infinium OmniExpress-24, and Illumina HumanCore Exome array) at participating centres. The *LRRK2* p.G2019S mutation was directly genotyped. Sanger sequence or KASPar verification of the *DNM3* rs2421947 variant was carried out on data subsets at study specific centres. Where the variant *DNM3* rs2421947 was imputed, only RSQ values above 0.85 were used, and only RSQ values above 0.75 were used for *VAMP4* rs11578699 (table-e1), as the RSQ quality of available data for variants *VAMP4* rs11578699 was lower. 1478 *LRRK2* p.G2019S carriers were identified across study cohorts. 2052 samples from patients with idiopathic PD from the Tracking Parkinson’s disease study passed quality control and had no identified genetic cause of disease (8). 96 patients with PD were excluded due to missing AAO data; therefore 1956 patients with PD and without *LRRK2* p.G2019S were included in further analysis. Summary statistics and patient data from the most recent PD GWAS included over 37.7K cases, 18.6K ‘proxy-cases’ and 1.4M controls. The most recent PD AAO GWAS included data from 28.6K patients (4,7).

#### Statistical analyses

Data from the studies were pooled. We calculated Hardy-Weinberg Equilibrium (HWE) for *VAMP4* rs11578699 and *DNM3* rs2421947 in p.G2019S carriers. QC HWE for iPD and controls is summarized in the recent large-scale PD GWAS. We assessed linkage disequilibrium (LD) measures RSQ and D’ between *DNM3* rs2421947 and *VAMP4* rs11578699 in different populations to evaluate the possibility that an extended haplotype block might explain the relationship between *DNM3* and *LRRK2* (table e-2). The allele based Fisher’s exact test was used to compare *DNM3* rs2421947 and *VAMP4* rs11578699 minor allele frequencies (MAF) between *LRRK2* p.G2019S heterozygotes of different ethnic backgrounds and with idiopathic PD (table 1). Student’s t-tests and ANOVA were used to assess differences in AAO by genotype (table 2). We then analyzed the effect of *DNM3* rs2421947 (figure 1) on AAO in idiopathic PD in the *Tracking Parkinson’s* dataset using linear regression of AAO, and Kaplan Meier survival analysis. Next, we analyzed patients with *LRRK2* p.G2019S of Ashkenazi Jewish, North African and European ethnicity using linear regression of AAO with available covariates of sex, ethnicity, relatedness and study centre of origin (figure 2). AAO was regressed by *DNM3* rs2421947 genotypes, and separately for rs11578699 *VAMP4* genotypes. We meta-analyzed *DNM3* rs2421947 data from all previously unpublished datasets using linear regression models (figure 2B). We pooled these data with previously published data and meta-analyzed again with the same methods (figure 2C and D). We then analyzed *VAMP4* rs11578699 impact on AAO with linear regression (figure 2E and F). Bonferroni or other corrections for multiple testing were not performed as this is a candidate gene-based study. Finally, we analyzed the impact of *VAMP4* rs11578699 on AAO in idiopathic PD carrying the *LRRK2* rs10878226 variant. *LRRK2* rs10878226 has also been implicated in PD risk (odds ratio [OR], 1.20; 95% confidence interval [CI], 1.08–1.33; p = 6.3×10^−4^, n=6129) (3).

**Figure 1.**
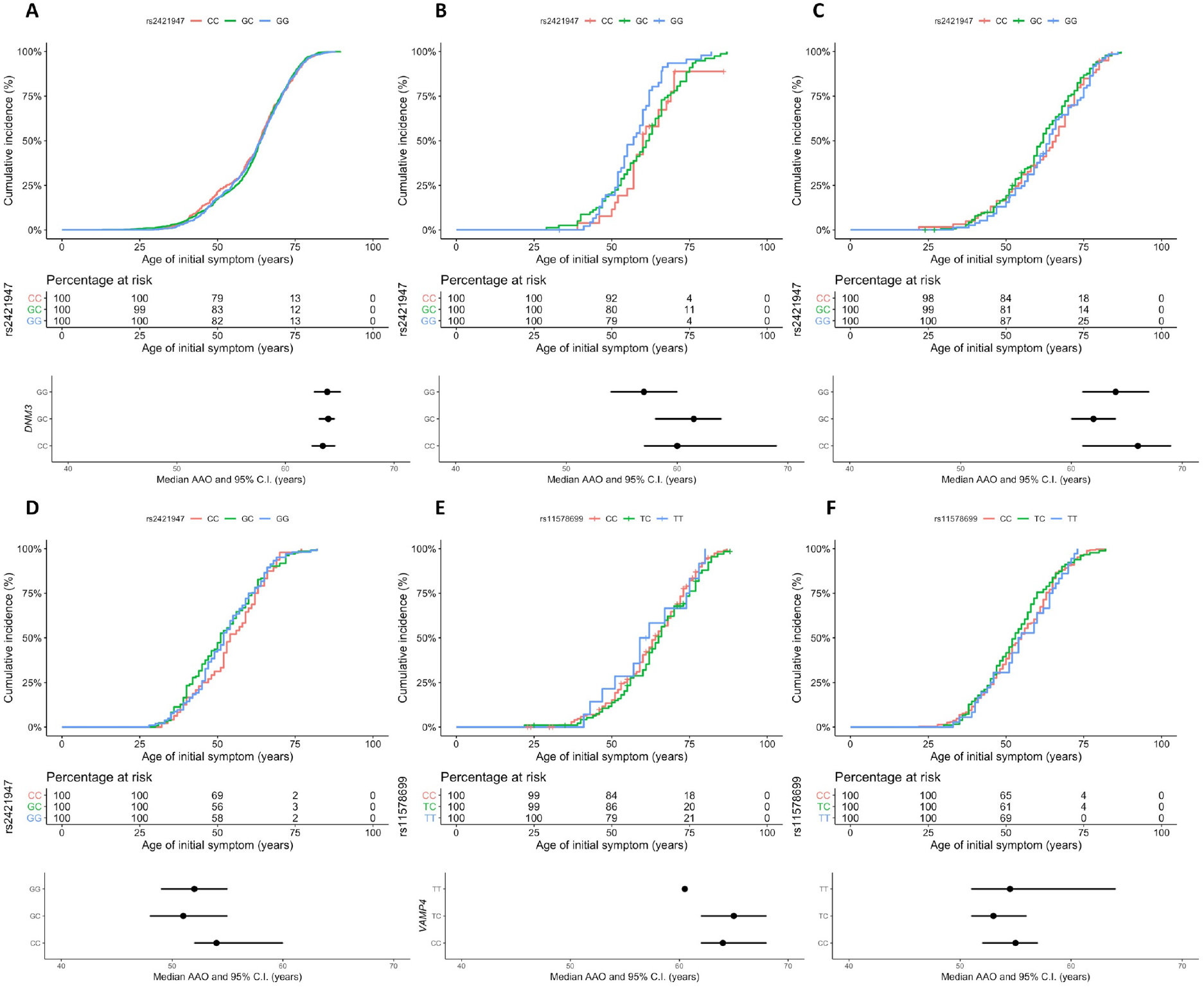
Survival analysis of AAO in *DNM3* rs2421947 (A-D) and *VAMP4* rs11578699 genotype groups (E-F) **A-C***DNM3* rs2421947 Kaplan Meier and Kaplan Meier median plots of **A** European idiopathic PD (n=1956); **B** Ashkenazi Jewish p.G2019S carriers (n=153) Kaplan Meier plot and median plot; **C** European p.G2019S carriers (n=286); **D** North African p.G2019S carriers (n=285). **E**-**F***VAMP4* rs2421947 Kaplan Meier and Kaplan Meier median plots **E** European and Ashkenazi Jewish p.G2019S carriers (n=302); **F** North African p.G2019S carriers (n=484).

**Figure 2.**
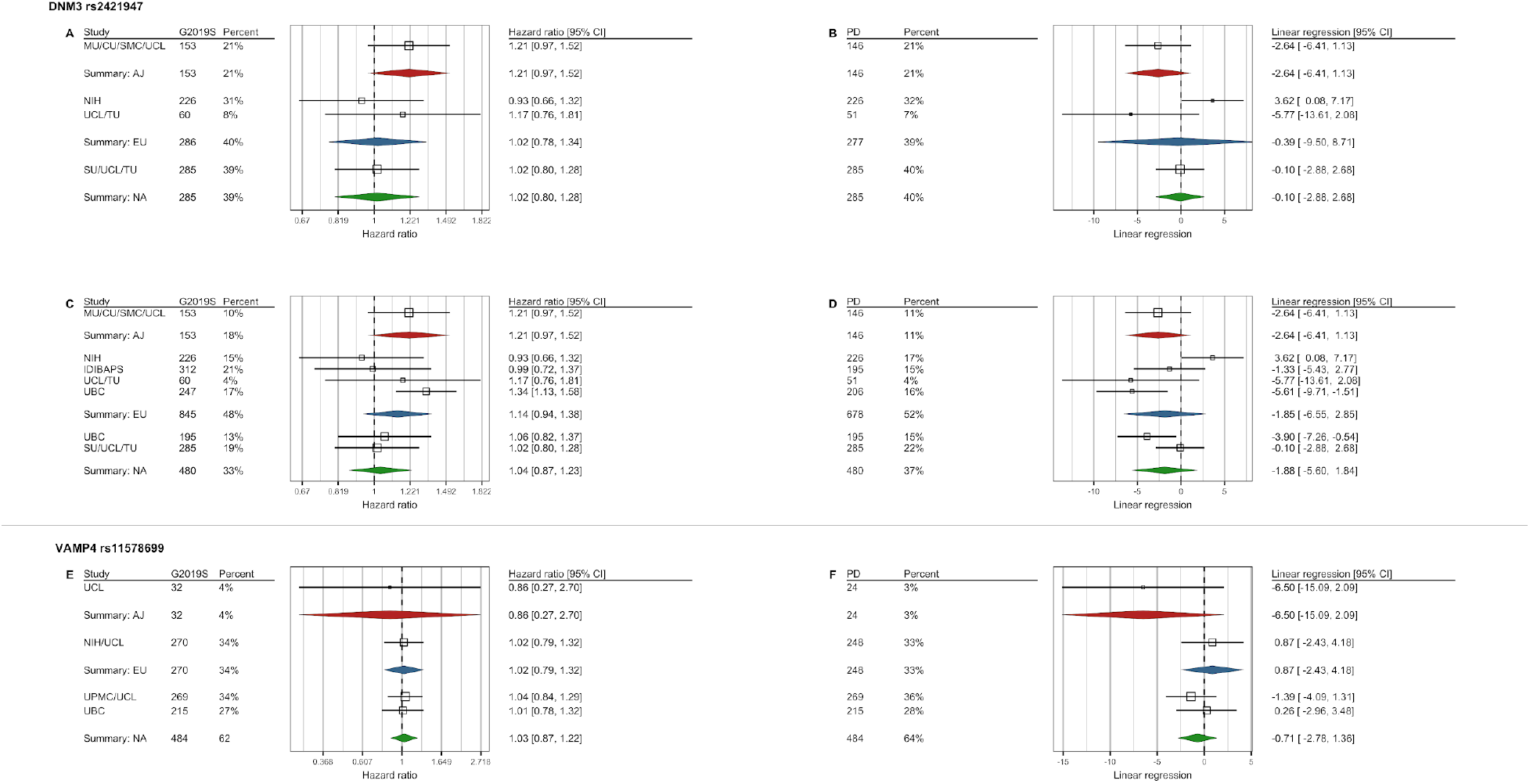
Forest plot of meta-analyses for *DNM3* rs2421947 and *VAMP4* rs11578699. (**A**, **C**, and **E**) Cox proportional hazards model meta-analyses *of LRRK*2 p.G2019S carriers, (**B**, **D**, and **F**) linear regression meta-analyses o*f LRRK*2 p.G2019S patients. **A** and **B**- meta-analyses of *DNM3* rs2421947 GG versus CG and CC genotypes of novel data from this manuscript from 724 *LRRK2* p.G2019S carriers; **C** and **D**- meta-analyses of *DNM3* rs2421947 GG versus CG and CC genotypes from 1478 *LRRK2* p.G2019S carriers including 754 previously published; **E** and **F**- *VAMP4* rs11578699 meta-analyses of CC versus TT and TC. Percentage contribution and numbers of individuals included in each analysis are indicated. Analyses are carried out on ethnicity subgroups: Ashkenazi Jewish (_Summary_: AJ), European (Summary: EU), and North African (Summary: NA).

**Table 1.**
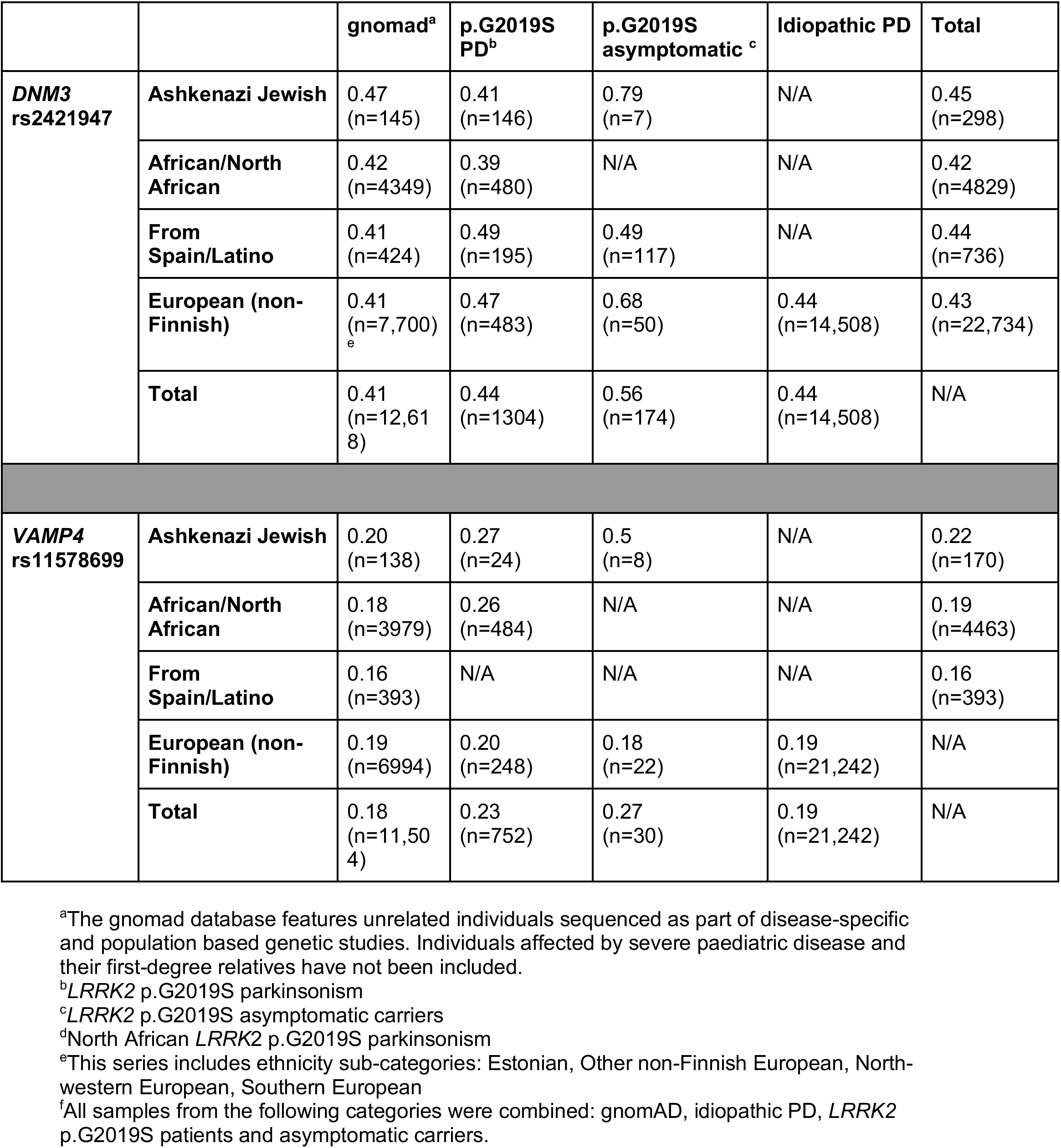
Minor allele frequency (MAF) for *DNM3* rs2421947 and *VAMP4* rs11578699 in PD cases and controls with and without p.G2019S.

|  |  | gnomad<sup>a</sup> | p.G2019S PD<sup>b</sup> | p.G2019S asymptomatic<sup>c</sup> | Idiopathic PD | Total |
| --- | --- | --- | --- | --- | --- | --- |
| ***DNM3* rs2421947** | **Ashkenazi Jewish** | 0.47<br>(n=145) | 0.41<br>(n=146) | 0.79<br>(n=7) | N/A | 0.45<br>(n=298) |
|  | **African/North African** | 0.42<br>(n=4349) | 0.39<br>(n=480) | N/A | N/A | 0.42<br>(n=4829) |
|  | **From Spain/Latino** | 0.41<br>(n=424) | 0.49<br>(n=195) | 0.49<br>(n=117) | N/A | 0.44<br>(n=736) |
|  | **European (non-Finnish)** | 0.41<br>(n=7,700)<br><sup>e</sup> | 0.47<br>(n=483) | 0.68<br>(n=50) | 0.44<br>(n=14,508) | 0.43<br>(n=22,734) |
|  | **Total** | 0.41<br>(n=12,618) | 0.44<br>(n=1304) | 0.56<br>(n=174) | 0.44<br>(n=14,508) | N/A |
| ***VAMP4* rs11578699** | **Ashkenazi Jewish** | 0.20<br>(n=138) | 0.27<br>(n=24) | 0.5<br>(n=8) | N/A | 0.22<br>(n=170) |
|  | **African/North African** | 0.18<br>(n=3979) | 0.26<br>(n=484) | N/A | N/A | 0.19<br>(n=4463) |
|  | **From Spain/Latino** | 0.16<br>(n=393) | N/A | N/A | N/A | 0.16<br>(n=393) |
|  | **European (non-Finnish)** | 0.19<br>(n=6994) | 0.20<br>(n=248) | 0.18<br>(n=22) | 0.19<br>(n=21,242) | N/A |
|  | **Total** | 0.18<br>(n=11,504) | 0.23<br>(n=752) | 0.27<br>(n=30) | 0.19<br>(n=21,242) | N/A |

<sup>a</sup>The gnomad database features unrelated individuals sequenced as part of disease-specific and population based genetic studies. Individuals affected by severe paediatric disease and their first-degree relatives have not been included.
<sup>b</sup>*LRRK2* p.G2019S parkinsonism
<sup>c</sup>*LRRK2* p.G2019S asymptomatic carriers
<sup>d</sup>North African *LRRK2* p.G2019S parkinsonism
<sup>e</sup>This series includes ethnicity sub-categories: Estonian, Other non-Finnish European, North-western European, Southern European
<sup>f</sup>All samples from the following categories were combined: gnomAD, idiopathic PD, *LRRK2* p.G2019S patients and asymptomatic carriers.

**Table 2.** Age at onset by genotype in *LRRK2* p.G2019S carriers in this study.

| ***DNM3* rs2421947** |  |  |  |  |  |  |  |
| --- | --- | --- | --- | --- | --- | --- | --- |
| Population series | Mean age at onset (sd) |  |  | ANOVA p value | GG vs CG p<sup>c</sup> | CG vs CC p<sup>c</sup> | GG vs CC p<sup>c</sup> |
|  | GG | CG | CC |  |  |  |  |
| Ashkenazi Jewish (n=146) | 57.20 (8.96) | 60.11 (12.21) | 58.41 (8.01) | 0.34 | 0.13 | 0.45 | 0.58 |
| North African (n=285) | 52.64 (11.21) | 51.95 (11.99) | 54.27 (11.05) | 0.49 | 0.65 | 0.23 | 0.40 |
| European<sup>a</sup> (n=277) | 63.26 (12.14) | 61.20 (12.45) | 62.17 (13.48) | 0.52 | 0.24 | 0.64 | 0.63 |
| Combined<sup>b</sup> (n=708) | 57.11 (12.02) | 57.44 (12.93) | 58.59 (12.30) | 0.55 | 0.76 | 0.37 | 0.28 |
| ***VAMP4* rs11578699** |  |  |  |  |  |  |  |
| Population series | CC | TC | TT | ANOVA p value | CC vs TC p<sup>c</sup> | TC vs TT p<sup>c</sup> | CC vs TT p<sup>c</sup> |
| Ashkenazi Jewish (n=24) | 63.92 (9.57) | 57.45 (11.93) | 57.00 (N/A) | 0.91 | 0.17 | N/A | N/A |
| North African (n=484) | 54.38 (11.89) | 53.35 (11.58) | 55.22 (11.48) | 0.55 | 0.36 | 0.38 | 0.68 |
| European<sup>a</sup> (n=248) | 62.44 (12.38) | 63.89 (12.21) | 61.33 (13.75) | 0.66 | 0.41 | 0.55 | 0.79 |
| Combined<sup>b</sup> (n=756) | 57.39 (12.61) | 56.41 (12.62) | 56.76 (12.10) | 0.60 | 0.32 | 0.85 | 0.73 |

Combined subgroup mean and standard deviation age at onset in years of *LRRK2* p.G2019S parkinsonism for *DNM3* rs2421947 and *VAMP4* rs11578699 genotypes; One-way ANOVA between *DNM3* rs2421947 and *VAMP4* rs11578699 genotypes; Student's t test between genotype
<sup>a</sup>The following population series subgroups from were combined: European unspecific, and British.
<sup>b</sup>The following population series subgroups from were combined: Ashkenazi Jewish, North African, European unspecific, and British.
<sup>c</sup>Uncorrected for multiple testing

#### Gene expression

Gene expression profiles of *DNM3* rs2421947 and *VAMP4* rs11578699 were assessed using publicly available Web-based resources: BRAINEACv.2 (www.braineac.org) (11); GTEx (www.gtexportal.org); and Allen Brain Atlas (www.brain-map.org) (12), and evaluated based on the significance level required for inclusion in colocalization (COLOC) analysis (appendix e-1).

## Results

### *DNM3* rs2421947 and V*AMP4* rs11578699 genotypes were in Hardy-Weinberg equilibrium in controls

There was no LD between *DNM3* rs2421947 and *VAMP4* rs11578699, as assessed through RSQ and D’, within or across population samples, indicating that these are independent variants (table e-2)

We investigated the possibility that ethnic variation in allele frequency might explain a variable effect of *DNM3* on affected *LRRK2* penetrance. Aside from the Norwegian cohort *DNM3* rs2421947 MAF (C) of 0.69 (vs. other Europeans p=0.00012), all *DNM3* rs2421947 MAF in the study were between 0.43-0.51 (table 1). There was no difference between Ashkenazi Jews and European frequencies (p=0.94); or between Ashkenazi Jews and North Africans (p=0.28); nor between Europeans and North Africans (p=0.54) at this locus. Publicly available allele frequencies (http://gnomad.broadinstitute.org) were similar (table 1). *VAMP4* rs11578699 MAF were between 0.15-0.30 in different populations.

We assessed possible association between these variants and p.G2019S genotype using the allele based Fisher’s exact test. There was no association between *DNM3* rs2421947 and p.G2019S genotype in all PD cases (p=0.42). There was no significant difference in *DNM3* allele frequencies between patients with PD and asymptomatic heterozygotes (asymptomatic p.G2019S heterozygotes were of Ashkenazi Jewish, North African, and European non-Jewish ethnicity), as assessed through Fisher’s Exact test (p=0.056). When the previous data was meta-analyzed with the current data there was a significant association between *DNM3* rs2421947 and PD affected status in p.G2019S heterozygotes (p=3.1×10^−5^) by the Fisher’s Exact test, although there were differences between populations (Table 1). Mean sampling age for p.G2019S heterozygotes without PD was 56 years across all ethnicities; 63 years in AJ subjects; and 56 years for Europeans.

In the most recent PD GWAS there was no genome-wide significant association between *DNM3* rs2421947 and PD by logistic regression, (p=0.0051). There was no association between *VAMP4* rs11578699 and *LRRK2* p.G2019S status in European PD (p=0.60) and no significant difference in *VAMP4* rs11578699 MAF between *LRRK2* p.G2019S patients and asymptomatic carriers by the Fisher’s Exact test (p=0.64).

We then assessed the relationship between AAO and *DNM3* rs2421947 and *VAMP4* rs11578699 (table 2) using t-tests and ANOVA. There was no effect of *DNM3* rs2421947 on AAO as assessed through ANOVA (p=0.55, F=0.59, df=2, n=708). When meta-analysed with data from previous cohorts there was a nominal effect of *DNM3* on AAO (2 years difference between G and C genotypes), (p=0.021, F=3.88, df=2, n=1304).

*DNM3* did not influence time to the development of disease in idiopathic PD for GG versus CC and CG carriers (beta = −0.02, p = 0.97, n =1956). This is consistent with the latest PD AAO GWAS, p=0.39, effect=−0.11, se=0.13. The Kaplan-Meier method was used for visualising risk across *DNM3* rs2421947 genotypes for idiopathic PD (Figure 1A), and LRRK2 p.G2019Scarriers (Figure 1B-D). The same method was also used to visualise risk for *VAMP4* rs11578699 genotypes in individuals with *LRRK2* p.G2019S (Figure 1E and F).

We carried out a multi-ethnic meta-analysis of newly contributed data of *DNM3* rs2421947 GG versus CC and CG genotypes against onset age in 724 *LRRK2* p.G2019S carriers, using a random effects model on disease-free survival. Linear regression meta-analysis on PD AAO was not significant (beta = −1.19, p = 0.55, n =708), though I^2^ heterogeneity was 86.9%, p < 0.01. The impact of GG versus CC and CG was not significant in sub-group analyses (figure 2). When our new data was pooled with the two previous studies and meta-analysed, linear regression meta-analysis of *LRRK2* p.G2019S AAO was not significant (beta = −2.21, p = 0.083, n = 1304). I^2^ total heterogeneity for linear regression meta-analysis was 82.3%, p<0.0001 (figure 2).

Meta-analysis of the impact of *VAMP4* rs11578699 (the genome-wide significant association in a recent large-scale Parkinson’s disease GWAS) on AAO i*n LRRK*2 p.G2019S parkinsonism was not significant for CC versus TT and TC genotypes (beta = −0.25, p=0.75, n=756).). I^2^ total heterogeneity was <0.001%, p=0.75 (figure 2).

We evaluated the possibility of ethnic specific effects using linear regression of AAO (figure 2). Though in each subgroup analysis confidence intervals overlapped zero, there appeared to be a trend towards earlier onset in G allele carriers of Ashkenazi Jewish ethnicity, consistent with previously reported data, and in the discovery data effects were strongest in Arab Berbers suggesting ethnic specific effects.

Using cox proportional survival analysis, we studied the discovery data and subsequent cohorts separately (GG versus CC and CG as described previously). There was a strong association between rs2421947 and PD AAO in discovery data (hazard ratio [HR] 1.59, 95% CI 1.28-1.97, p<0.001). This was also present through AAO linear regression analysis (beta = −4.93, p = 0.00019). The association was absent in our replication data (hazard ratio [HR] 1.09, 95% CI 0.95-1.26, p = 0.22).

We meta-analyzed the samples studied in this paper with the discovery data (Figure 2).

Finally, we investigated the possibility that might be an interaction between the GWAS defined common *LRRK2* risk allele (rs10878226) and *VAMP4*. We analyzed 4882 cases with PD carrying the *LRRK2* risk variant minor allele using linear regression of AAO and plotting of the Kaplan Meier curve, as shown in Figure 3. TT versus TC and CC *VAMP4* rs11578699 genotype was nominally significantly associated with AAO risk in this cohort (beta=1.68, se=0.81, p=0.037, n=4882*). VAMP4* rs11578699 was not significantly associated with AAO risk in idiopathic PD cases without the *LRRK2* rs10878226 risk variant (beta=-0.28, se=0.48, p=0.56, n=14970). Far fewer patients with idiopathic PD had been genotyped for *DNM3* rs2421947 in the PD GWAS cohort (table 1), meaning we were not able to assess interaction between *DNM3* and the *LRRK2* risk allele.

**Figure 3.**
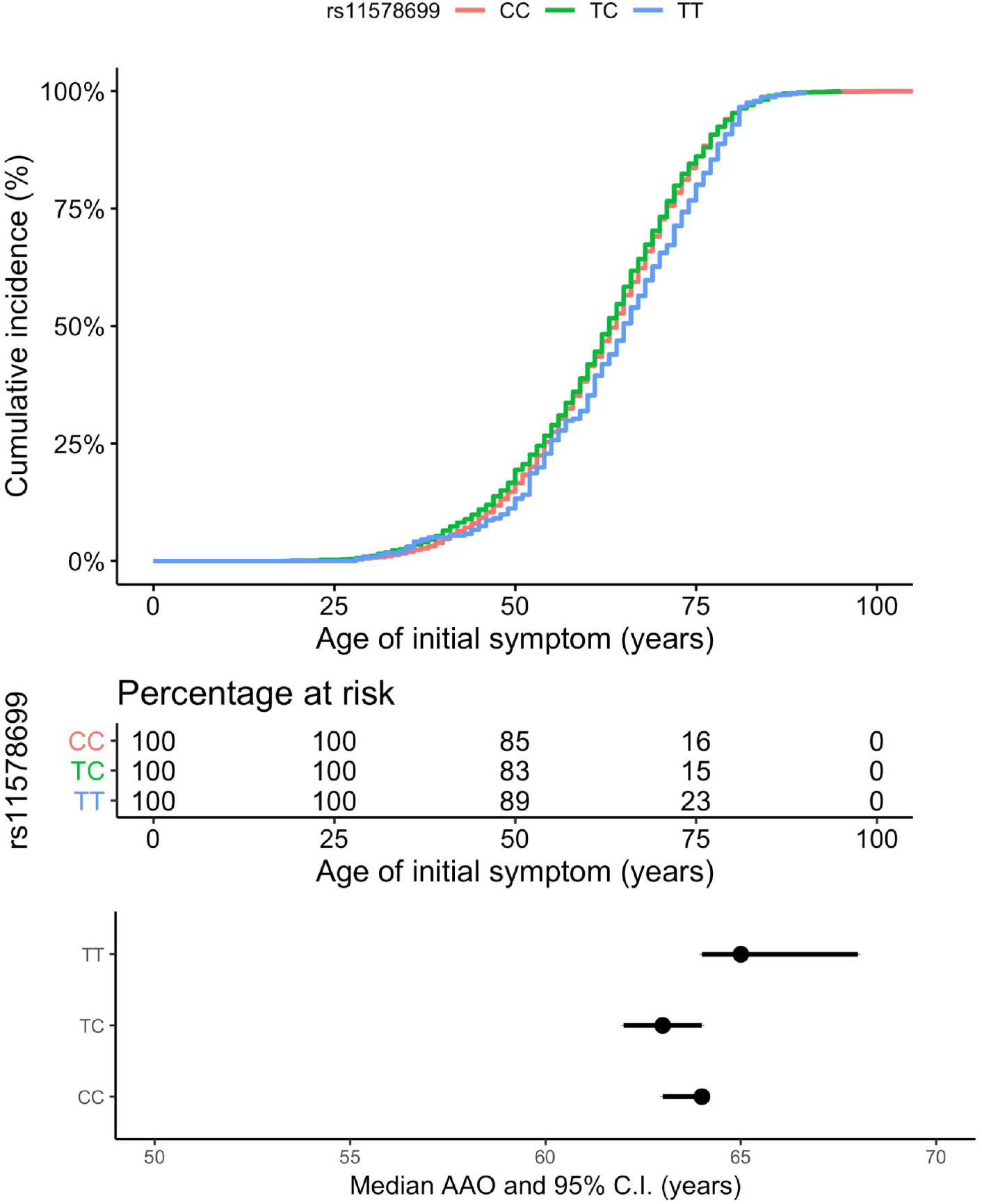
Kaplan Meier analysis by *VAMP4* rs11578699 genotype in PD cases carrying *LRRK2* risk variant (rs10878226)

We then reviewed *VAMP4* and *DNM3* human brain gene expression and co-expression data from the BRAINEACv2^23^ (11), GTEx, and Allen Atlas databases (12) (figure e-3)*. VAMP4* rs11578699 is an eQTL for *VAMP4* expression: NES: 0.29, p = 3.5×10^−9^ (cerebellum, GTEx database); NES: 0.32, p = 6.5×10^−7^ (cerebellar hemisphere, GTEx database). *DNM3* rs2421947 did not appear to be an eQTL for *DNM3* expression in the brain in the GTex database, though *VAMP4* and *DNM3* gene expression in brain were high (figure e-3).

## Discussion

We have analyzed the effect of *DNM3* rs2421947 on AAO in *LRRK2* p.G2019S parkinsonism. We have not replicated the association between *DNM3* and *LRRK2* p.G2019S AAO in this study or in meta-analysis with all available data. In the original discovery analysis, there was a difference of 12.5 years between *DNM3* rs2421947 GG and CC genotypes (meta-analysis HR 1·61, 95% CI 1·15–2·27, p=0·02). In our data the AAO difference between GG and CC genotypes was 1.4 years, which was not significant. Using AAO regression, meta-analysis of independent sample series in newly genotyped samples did not identify a significant difference in AAO between genotypes (beta = −1.19, p = 0.55, n =708). Similarly, when discovery and replication data was analyzed together there was no significant effect of *DNM3* rs2421947 on AAO (beta = −2.21, p = 0.083, n = 1304). However, there was significant heterogeneity in replication and combined *DNM3* regression meta-analyses.

In “sporadic” PD, consistent with the recent AAO GWAS (4), our study indicated that *DNM3 rs2421947* does not affect AAO in non-p.G2019S disease.

We did not identify linkage disequilibrium between DNM3 and VAMP4, nor an independent effect of VAMP4 on LRRK2 penetrance. Analysis of carriers of a common *LRRK2* risk SNP provided nominally significant support for a potential interaction between *VAMP4* and *LRRK2,* which requires replication in larger sample sizes. Gene expression analysis indicated that *VAMP4* rs11578699 is an eQTL for *VAMP4* expression.

One possibility for the lack of replication of *DNM3* modification of p.G2019S AAO is that heterogeneity in the sample may limit the observed effect in the meta-analysis; this has previously led to non-replication in independent samples in genome-wide association studies (13) (14). Significant heterogeneity was observed in the linear regression meta-analysis of replication and combined data in this study, which may be explained by ethnicity effects. The impact of *DNM3* may vary between ethnicities or be population specific, and the strongest effects were seen in Ashkenazi Jewish p.G2019S carriers although not reaching significance (Figure 2). Further studies in large numbers of Ashkenazi Jewish and Arab patients are needed.

Due to its role in innate immunity *LRRK2* may have been under different selective pressures in human evolution, relating to differences in environmental pathogens. Interestingly, *LRRK2* p.G2019S penetrance varies across ethnic groups. Hentati and colleagues (15) have reported a later AAO in *LRRK2* p.G2019S carriers from Norway as compared to Tunisian patients. Lee and colleagues (16) indicated later AAO penetrance for Ashkenazi Jewish PD p.G2019S carriers compared to non-Jewish carriers through plotting of estimated cumulative risk, though this was not significant; penetrance was 42.5% (95% CI: 26.3-65.8%) in non-Ashkenazi Jewish relatives compared to 25% (95% CI: 16.7%-34.2%) in Ashkenazi Jewish heterozygous relatives. These data imply that there may be potential protective factors in the relatively homogeneous Norwegian and Ashkenazi Jewish population and that ethnic and population effects are likely to be very important in analyzing this variant. Our analysis and comparison of background allele frequencies in this study suggest that this is unlikely to primarily relate to *VAMP4* and *DNM*3. However, there may be other rare variant effects that will emerge with fine mapping of this region.

The *LRRK2* common risk variant analysis is of interest; the variant is not a proxy for p.G2019S in Europeans, so this represents an independent marker of PD risk. Around a quarter (24.6%) of sporadic PD cases carry the common *LRRK2* risk variant rs10878226, which is associated with PD (combined odds ratio [OR], 1.20, 95% CI, 1.08-1.33, p=6.3×10^−4^, n=6129) (3). Our common *LRRK2* variant analysis indicates a possible interaction between common variation in *LRRK2* and *VAMP4,* which requires further study. *VAMP4* and *LRRK2* are both involved in synaptic vesicle dynamics, which has relevance to the etiology of PD. Underscoring this possibility, analysis of *VAMP4* and *DNM3* expression indicated that both are highly expressed in the brain, though these are not in LD and likely represent independent signals.

Modifiers of the penetrance of LRRK2 p.G20129S are likely to be therapeutic targets and may be important in genetic counselling. Our large study, aggregating new and previously published data has indicated significant heterogeneity across studies, but not provided robust replication of an interaction between *DNM3* and *LRRK2* p.G2019S. Further genome-wide studies in different populations are needed, to resolve the determinants of the variable penetrance seen in individuals with p.G2019S.

## Acknowledgement

We would like to thank all of the subjects who donated their time and biological samples to be part of this study.

This work was supported by Parkinson’s UK, University College London (UCL), and the Medical Research Council (MRC). Research ethics approval was provided by West of Scotland Research Ethics Service (reference 11/AL/0163) and multiple regional ethics committees. The study is registered with ClinicalTrials.gov under the identifier NCT02881099.

We would also like to thank all members of the International Parkinson Disease Genomics Consortium (IPDGC). For a complete overview of IPDGC members, acknowledgements and funding, please see: http://pdgenetics.org/partners. This work was supported in part by the Intramural Research Programs of the National Institute of Neurological Disorders and Stroke (NINDS) and the National Institute on Aging (NIA).

The study was funded by the Medical Research Council and the NIHR Rare Diseases Translational Research Consortium.

Funding for the PFP study was provided by University College London (UCL), Parkinson’s UK and the Medical Research Council. Research ethics approval was provided by the London Camden and King’s Cross ethics committee (reference 15/LO/0097) and the Health Research Authority (HRA). The study is registered with ClinicalTrials.gov under the identifier NCT02760108.

The authors thank the PD patients and unaffected relatives who participated in this study.

## Supplementary material

**Table e-1.** Imputation quality per subgroup.

|  |  |
| --- | --- |
| **RSQ imputation score** |  |
| **iPD *DNM3* rs2421947** |  |
| >0.99 | 100% |
| **p.G2019S carriers *DNM3* rs2421947** |  |
| 1.00 | 70% |
| >0.99 | 73% |
| >0.90 | 98% |
| **p.G2019S carriers *VAMP4* rs11578699<sup>1</sup>** |  |
| 1.00 | 79% |
| >0.98 | 86% |
| >0.75 | 100% |
| **iPD *VAMP4* rs11578699** |  |
| Previously published PD GWAS data<sup>1</sup> | 100% |

## appendix e-1 Supplementary methods

For the meta-analyses data was divided into the same ethnicity subgroups (Ashkenazi Jewish, North African, and European), and study centre specific groups. Where there were <60 datapoints from a particular study centre for *DNM3* rs2421947, data was pooled, and study centre of origin was used as a covariate. Cox proportional hazards model analyses and linear regressions for p.G2019S carriers were carried out on GG versus CC and CG genotypes, as exploratory data analysis indicated a dominant protective effect of the minor C allele. For the *VAMP4* rs11578699 variant these survival methods were carried out on CC versus TC and TT due to the smaller number of TT genotypes available. Family relatedness, sex, and principal components of ancestry were used as covariates, where available, in each analysis.

A random-effects model was used in meta-analyses. Natural logs of hazard ratios for GG versus CC and CG genotypes were taken for *DNM3* rs2421947, and for CC versus TC and TT for *VAMP4* rs11578699. The log of standard errors was calculated using outputted hazard ratio confidence intervals, using the Taylor expansion with leading term (also known as the delta method) (17).

RSQ and D’ were calculated in PLINK 1.9 (18). The allele based Fisher’s Exact test, ANOVA, Student’s t tests, and linear regressions (PD onset age regressed on *DNM3* rs2421947 genotype, and separately *VAMP4* rs11578699 genotype) were performed in base R RStudio (19). Cox proportional hazards survival analysis and Kaplan Meier curves were carried out using the Terry Therneau “Survival” package (20) in RStudio (19). Time-to-event in survival analyses was the self-reported age at which a participant developed motor symptoms of PD. For p.G2019S carriers without PD time-to-event was right-censored from sampling age. Meta-analysis was carried out in metafor (21) and metaviz R packages (22).

We supplemented the analysis of *LRRK2* p.G2019S by analysing time-to-event (by *DNM3* rs2421947 and *VAMP4* rs11578699) genotypes in PD cases carrying the *LRRK2* risk variant rs10878226 (4). We identified 4882 PD cases heterozygous or homozygous for the minor risk allele from the 21,242 PD cases in IPDGC datasets. Linear regression of AAO, Kaplan Meier and Cox proportional hazards analyses were then performed on *VAMP4* rs11578699 genotypes against age at onset. These methods were carried out on TT versus TC and CC genotypes as exploratory data analysis indicated an effect of the homozygous T allele on AAO. Covariates of sex, ethnicity (PC1-PC10) and dataset were used. We carried out the same analysis on 14970 PD cases not carrying the *LRRK2* rs10878226 variant.

## Gene Expression

The BRAINEACv2 database hosts data on brain tissues (frontal cortex, temporal cortex, parietal cortex, occipital cortex, hippocampus, thalamus, putamen, substantia nigra, medulla, cerebellum, and white matter) from 134 healthy controls. The GTEx database consists of 8555 samples from 53 tissues (including 13 brain regions) from 544 donors. The Allen Human Brain Atlas database microarray data is from 8 neuropathologically normal individuals of varying ethnicity, and covers ~150 brain regions.

---

**table e-2.** Population specific D’ and RSQ values between *DNM3* rs2421947 and *VAMP4* rs11578699.

|  | Controls | PD cases | p.G2019S PD cases |
| --- | --- | --- | --- |
| **D'** |  |  |  |
| **Ashkenazi Jewish** | N/A | N/A | 0.087 (n=29) |
| **African/North African** | 0.089 (n=1322)<sup>a</sup> | N/A | 0.061 (n=482) |
| **From Spain/Latino** | 0.30 (n=214)<sup>b</sup> | N/A | N/A |
| **European non-Finnish** | 0.46 (n=594)<sup>c</sup> | 0.21 (n=14508) | 0.27 (n=186) |
| **Total** | 0.22 (n=2130) | 0.21 (n=14508) | 0.021 (n=697) |
| **RSQ** |  |  |  |
| **Ashkenazi Jewish** | N/A | N/A | 0.0035 (n=29) |
| **African/North African** | 0.0013 (n=1322)<sup>a</sup> | N/A | 0.0022 (n=482) |
| **From Spain/Latino** | 0.037 (n=214)<sup>b</sup> | N/A | N/A |
| **European non-Finnish** | 0.039 (n=594)<sup>c</sup> | 0.0086 (n=14508) | 0.017 (n=186) |
| **Total** | 0.0085 (n=2130) | 0.0086 (n=14508) | 0.000098 (n=697) |

<sup>a</sup>All (AFR) Populations: (YRI, LWK, GWD, MSL, ESN, ASW, ACB)
<sup>b</sup>Selected (EUR) population: (IBS)
<sup>c</sup>Selected (EUR) population: (CEU, GBR, TSI)

## Supplementary results

### Cox proportional hazard analysis

*DNM3* did not influence time to the development of disease in idiopathic PD by the Cox proportional hazards model (hazard ratio [HR] 0.98, 95% CI 0.89-1.07, p = 0·60, n=1956) for GG versus CC and CG carriers. *DNM3* rs2421947 did not significantly affect the age associated hazards of developing p.G2019S parkinsonism through random effects meta-analysis in our previously unpublished data (hazard ratio [HR] 1.09, 95% CI 0.95-1.25, p = 0.20, n=724), as shown in Figure 2. I^2^ heterogeneity was <0.01% (p = 0.42). When our new data was pooled with the two previous studies and meta-analysed, there was a nominally significant effect of *DNM3* rs2421947 on AAO in *LRRK2* p.G2019S parkinsonism ([HR] 1.14, 95% CI, 1.02-1.27, p = 0.025, n = 1478). I^2^ total heterogeneity was 22.8%, p=0.37.

*VAMP4* rs11578699 was nominally associated with disease AAO in idiopathic PD cases carrying the *LRRK2* risk variant rs10878226 (TT versus CC and TC (hazard ratio [HR] 0.86, 95% CI 0.76-0.98, p=0.023), and not associated with AAO in idiopathic PD cases not carrying the *LRRK2* risk variant rs10878226 (hazard ratio [HR] 1.02, 95% CI 0.94-1.10, p=0.47).

**figure e-1.**
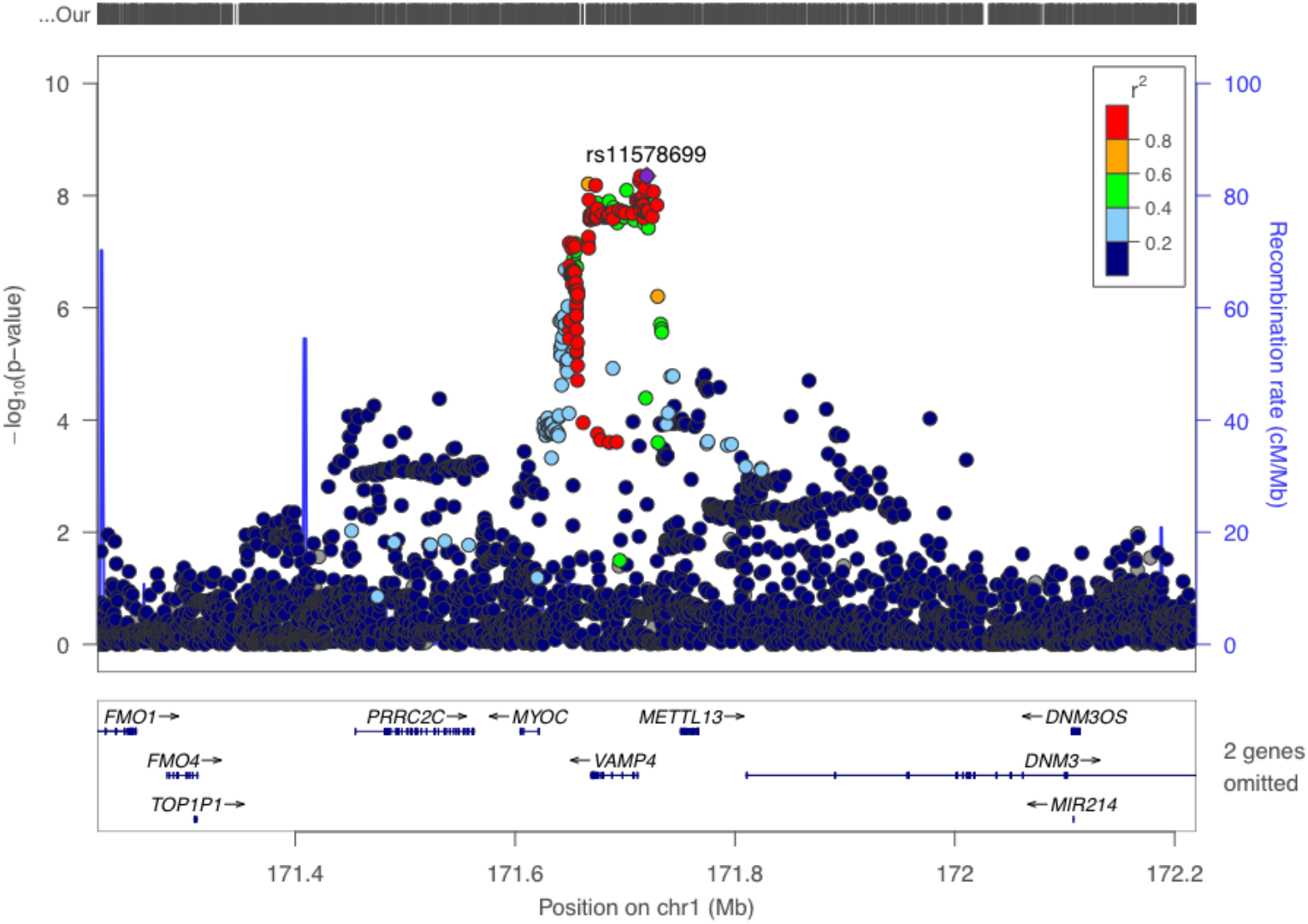
Regional association plot of PD cases versus control from the largest PD GWAS, identifying rs11578699 as the lead SNP.

**table e-3.** Population series mean and standard deviation age at onset in years of p.G2019S PD cases for rs2421947 genotypes; One-way ANOVA between rs2421947 genotypes; Student’s t test between rs2421947 genotypes.

| Population series | Mean age at onset (sd) |  |  | ANOVA p value | t test GG versus CG p value<sup>b</sup> | t test CG versus CC p value<sup>b</sup> | t test GG versus CC p value<sup>b</sup> |
| --- | --- | --- | --- | --- | --- | --- | --- |
|  | GG | CG | CC |  |  |  |  |
| **European unspecific (n=239)<sup>a</sup>** | 65.35 (11.15) | 62.19 (12.19) | 62.51 (14.00) | 0.23 | 0.075 | 0.88 | 0.24 |
| **American (n=87)** | 56.30 (7.53) | 61.76 (10.14) | 64.55 (10.98) | 0.16 | 0.061 | 0.44 | 0.058 |
| **French (n=98)** | 52.08 (9.96) | 50.93 (13.48) | 49.94 (10.51) | 0.81 | 0.66 | 0.77 | 0.49 |
| **Norwegian (n=18)** | 44.00 (N/A) | 72.55 (15.12) | 61.67 (20.81) | 0.21 | Insufficient t count<sup>c</sup> | 0.29 | Insufficient t count<sup>c</sup> |
| **Spanish (n=195)** | 58.70 (14.01) | 59.93 (12.59) | 60.24 (12.78) | 0.81 | 0.60 | 0.89 | 0.56 |
| **British (n=36)** | 49.68 (9.54) | 53.78 (11.20) | 60.00 (9.99) | 0.13 | 0.32 | 0.18 | 0.042 |
| **Ashkenazi Jewish (n=146)** | 57.20 (8.96) | 60.11 (12.21) | 58.41 (8.01) | 0.34 | 0.13 | 0.45 | 0.58 |
| **North African (n=480)** | 53.02 (11.89) | 54.16 (11.41) | 55.75 (12.11) | 0.23 | 0.33 | 0.32 | 0.10 |

<sup>a</sup>Samples defined as from a non-specific European background (through PCA or self-identification during clinician interview).
<sup>b</sup>Uncorrected for multiple testing
<sup>c</sup>There are insufficient p.G2019S carriers in these categories for the t test to be calculated.

**figure e-2.**
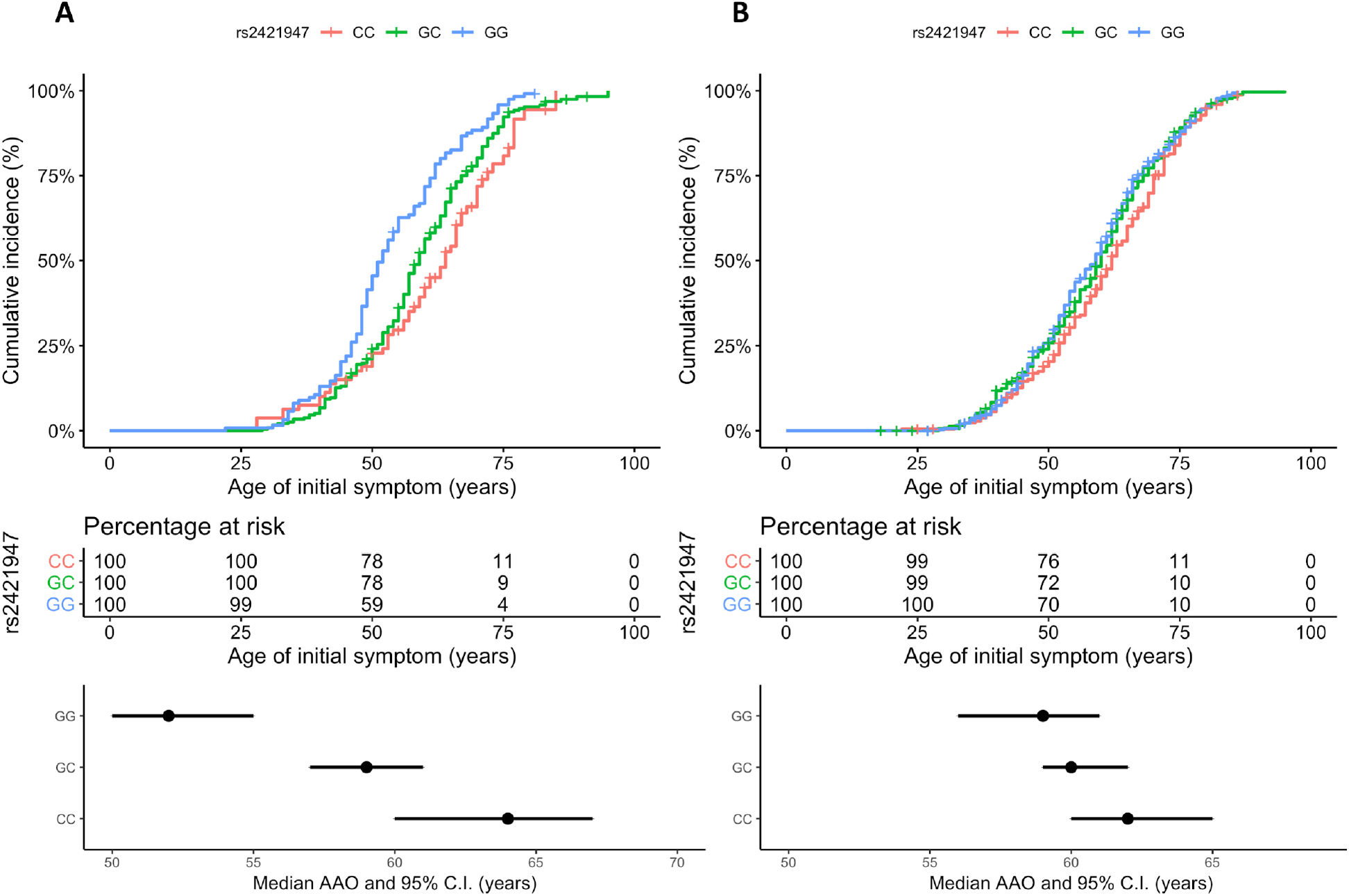
Kaplan Meier plots by rs2421947 genotypes-comparison of discovery data and subsequent cohorts. (A) Discovery data (n = 440) p.G2019S carriers Kaplan Meier plot and median plot (B) Subsequent cohorts (n = 1038) p.G2019S carriers Kaplan Meier plot and median plot.

**figure e-3.**
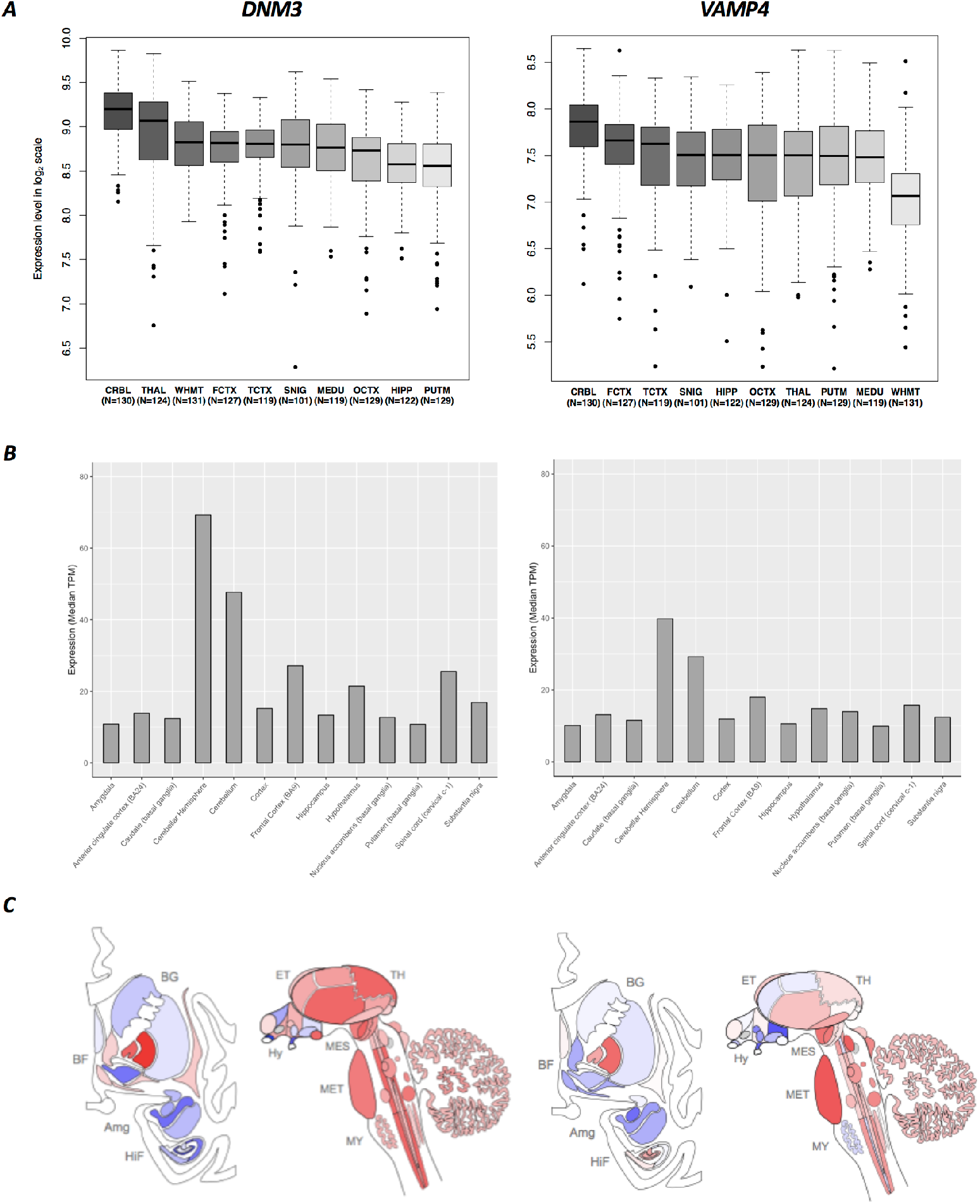
Regional variation in the brain of *DNM3* and *VAMP4*. DNM3 and VAMP4 brain expression in: **A.** BRAINEAC database – CRBL = Cerebellum, PUTM = Putamen, HIPP = Hippocampus, TCTX = Temporal cortex, OCTX = Occipital cortex, FCTX = Frontal cortex, THAL = Thalamus, SNIG = Substantia nigra, MEDU = Medulla, WHMT = White matter; **B.** GTEx database; **C.** Allen Atlas database (European subjects) – Amg = Amygdala, BF = Basal Forebrain, BG = Basal Ganglia, CAU = Caudate, ET = Epithalamus, HiF = Hippocampal Formation, Hy = Hypothalamus, MES = Mesencephalon, MET = Metencephalon, MY = Myelencephalon, PUT = Putamen, TH = Thalamus. Image credit: Allen Institute.

